# Species-specific CD4^+^ T cells Enable Prediction of Mucosal Immune Phenotypes from Microbiota Composition

**DOI:** 10.1101/2022.08.13.503851

**Authors:** Matthew P. Spindler, Ilaria Mogno, Prerna Suri, Graham J. Britton, Jeremiah J. Faith

## Abstract

How bacterial strains within a complex human microbiota collectively shape intestinal T cell homeostasis is not well-understood. Methods that quickly identify effector strains or species that drive specific mucosal T cell phenotypes are needed to define general principles for how the microbiota modulates host immunity. We colonize germ-free mice with defined communities of cultured strains and profile antigen-specific responses directed towards individual strains *ex vivo*. We find that lamina propria T cells are specific to bacterial strains at the species level and can discriminate between strains of the same species. *Ex vivo* restimulations consistently identify the strains within complex communities that induce Th17 responses *in vivo* providing the potential to shape baseline immune tone via community composition. Using an adoptive transfer model of colitis, we find that lamina propria T cells respond to different bacterial strains in conditions of inflammation versus homeostasis. Collectively, our approach represents a novel method for efficiently predicting the relative impact of individual bacterial strains within a complex community and for parsing microbiota-dependent phenotypes into component fractions.

**SIGNIFICANCE:** Determining the mechanisms by which the gut microbiome modulates the host immune system has translational potential for treating or preventing immune mediated disease. A key challenge is identifying the immunogenic bacterial strains in the setting of a complex microbiota. We use a combination of anaerobic culturing, *in vitro* T cell assays, gnotobiotic mouse models, and *ex vivo* T cell restimulations to explore the influence of species and strain diversity on the specificity of mucosal T cells. Our approach efficiently predicts the relative impact of individual bacterial strains within a complex community and can be used to parse microbiota-dependent phenotypes into component fractions.

## INTRODUCTION

The microbiota plays a fundamental role in the education of the immune system (Belkaid and Hand, 2014). Despite known associations between the microbiota and a range of inflammatory diseases, we have limited understanding of the specificity at which antibodies, T cells, and the innate immune system recognize the complexity of the microbiome. In addition, our knowledge of which bacterial strains influence different immune populations is limited to a few characteristic examples (e.g., Atarashi et al., 2011; Britton et al., 2020; Faith et al., 2014; Viladomiu et al., 2017). Therefore, we remain limited in our ability to harness microbial communities to alter host immune function and treat immune mediated diseases.

In the context of T cells, the field has established that gut microbiotas differentially modulate Th1, Th2 and Th17 cells (Atarashi et al., 2017; Britton et al., 2019; Ivanov et al., 2008; Yang et al., 2014). However, identifying the bacterial strains within a complex human microbiota that shape intestinal T cell phenotypes has proven challenging. Besides several limited examples, we do not understand the molecular mechanisms and the microbial signals that lead to specific T cell responses in the gut (Ansaldo et al., 2019; Ivanov et al., 2009; Nagashima et al., 2022; Tan et al., 2016). Efforts to parse microbiota-dependent immune phenotypes have either used time-intensive methods or relied upon trial-and-error *in vivo* experimentation (Atarashi et al., 2015, 2011; Britton et al., 2020; Britton and Faith, 2021; Faith et al., 2014; Geva-Zatorsky et al., 2017; Surana and Kasper, 2017; Viladomiu et al., 2017). Furthermore, given the broad compositional differences within disease-associated human microbiotas (Contijoch et al., 2019; Gevers et al., 2014), it is not known if the recognition of microbiota-derived strains by the host immune system depends on the host environment, such as the presence or absence of inflammation. Methods that quickly identify the effector strains driving mucosal T cell phenotypes are needed to define more general principles for how the microbiota modulates host immunity in both healthy and disease-states.

Here, we address three fundamental questions regarding how mucosal T cells interact with gut microbiota-derived bacterial strains. 1) What is the specificity of microbially-primed T cells? 2) How do microbial strains in a complex community collectively influence mucosal immune tone? And 3) Does host health status impact the T cell response towards the gut microbiota? To address these questions, we leverage a combination of *in vitro, in vivo*, and *ex vivo* model systems and use individual bacterial strains or collections of cultured strains to prime the host immune system. We subsequently read out the specificity of the T cell response towards individual strains using recall assays such as ELISpot. We find across both *in vitro* and *in vivo* model systems that Th17 cells are specific to bacteria at the species level and can partially differentiate between strains of the same species. Furthermore, we use our *ex vivo* T cell restimulation approach to predict the relative *in vivo* impact of individual bacterial strains within larger more complex communities. These methods repeatedly and efficiently identify the effector strains within complex culture communities that induce Th17 responses *in vivo*. Remarkably, we find the *ex vivo* restimulation approach for each individual strain is additive, allowing us to predict the mucosal T cell phenotype of random communities as a function of the immunostimulatory capacity of individual strains, raising the potential of engineering desired host immune phenotypes using selective microbiota-derived bacterial strains. Finally, using an adoptive transfer model of colitis, we show that lamina propria T cells respond to different bacterial strains in homeostatic versus inflammatory settings.

## RESULTS

### Specificity of *in vitro* primed T cells towards gut-derived bacterial strains

To gain insights into the T cell specificities towards the microbiota, we first modeled *in vitro* interactions between T cells and antigens from microbiota-derived bacterial strains. Using a recently established approach (Gao et al., 2020), we loaded 20,000 CD11c^+^ dendritic cells (DCs) with antigen from lysates of one of three different human gut microbiota-derived strains: *Escherichia coli strain E, Ruminococcus gnavus strain A*, and *Bacteroides ovatus strain C* (Supplementary Tables 1 and 2). We then primed 100,000 naïve CD4^+^ T cells by coculturing them with antigen-loaded DCs to generate oligoclonal T-cell cultures educated towards a single bacterial strain (Figure 1A). To broadly examine the relative specificities towards a diverse set of microbiota-derived antigens, we restimulated these 3 different *in vitro* educated T cell cultures using irradiated B cells to present antigens from a library of 45 bacterial strain lysates – 4 Actinobacteria, 15 Bacteroides, 19 Firmicutes, 7 Proteobacteria (including multiple strains of the same species) (Figure 1B, Supplementary Tables 1 and 3). T cells primed with either an *E. coli* strain lysate, an *R. gnavus* strain lysate, or a *B. ovatus* strain lysate elicited a diverse range of IFNγ and IL-17A responses when restimulated with the 45-strain antigen library (*E. coli* p < 1×10^−10^, ANOVA, Figures 1C, S1A and S1B). For both *E. coli* and *B. ovatus*, we found a strong correlation between IL-17A and IFNγ recall responses towards each antigen in the library (*E. coli* R^2^ = 0.91, p < 1×10^−10^, *B. ovatus* R^2^ = 0.66, p < 1×10^−10^, Figure S1C), while these cytokine responses were less correlated for *R. gnavus* (R^2^ = 0.07, p = 0.04, Figure S1C). Overall, IFNγ had a larger dynamic range across all three priming conditions (Figure S1C) and therefore we concentrated the remainder of our *in vitro* analyses on IFNγ responses to maximize the sensitivity of the assay.

**Figure 1.**
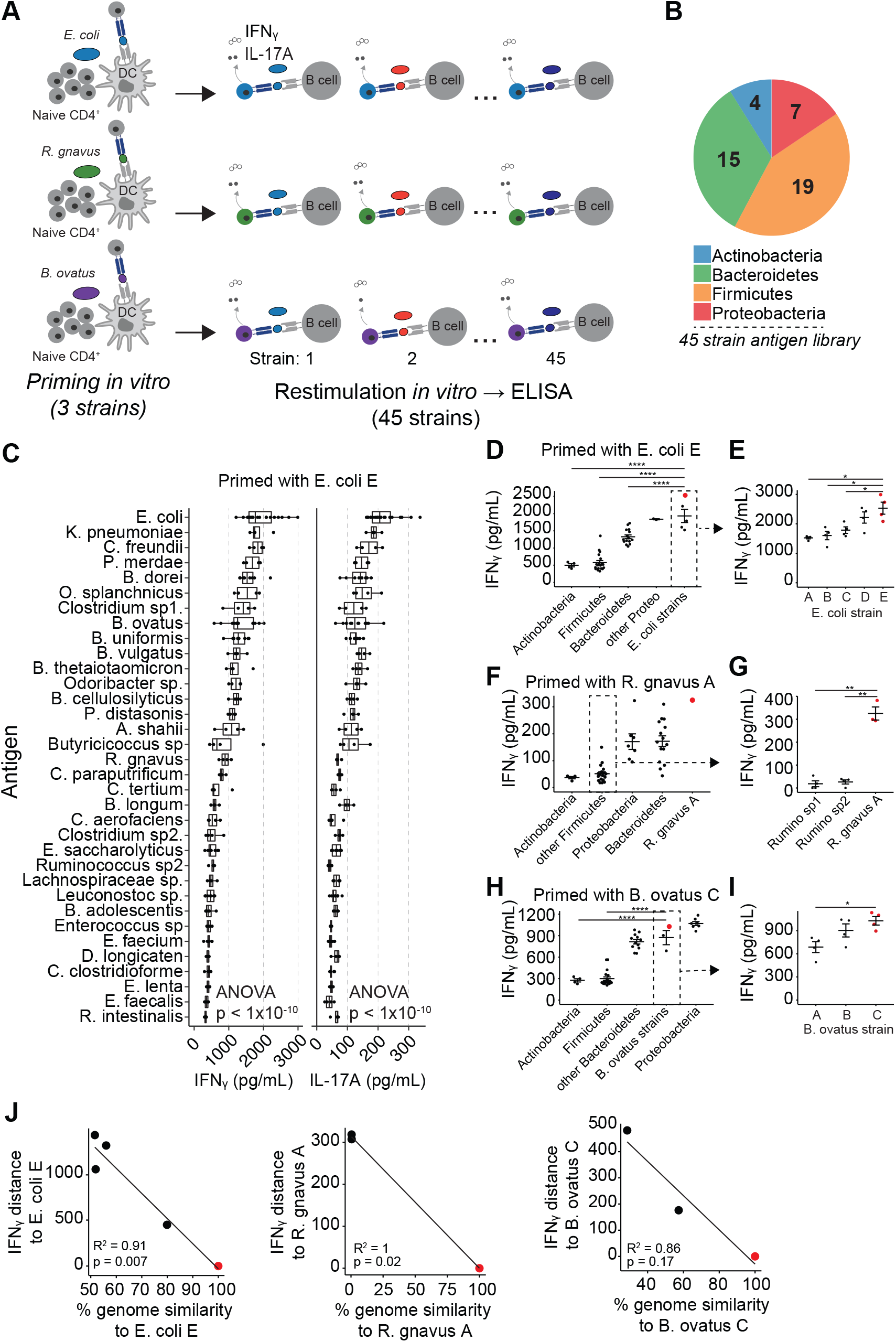
T cells primed *in vitro* are specific towards gut-derived bacterial strains. (A) Schematic of experimental design. 100,000 CD4^+^ CD44^-^ CD62L^+^ naïve T cells were co-cultured with 20,000 CD11c^+^ dendritic cells loaded with a lysate from a microbiota-derived bacterial strain for 7 days. T cells were restimulated using antigens from 45 unique bacterial strains loaded on CD19^+^ B cells. IFNγ and IL-17A were measured from the supernatant by ELISA 48 hours following restimulation. (B) Bacterial antigen library consists of 45 strains from the four major phyla in the human gut microbiota. (C) IFNγ and IL-17A levels as measured by ELISA from T cells following restimulation with the indicated bacterial strains. T cells were primed with *E. coli E*. (D) IFNγ as measured by ELISA from T cells primed with *E. coli E* and restimulated with bacterial strains from the indicated taxa. Student’s t-test. **** p <0.0001. (E) IFNγ from T cells primed with *E. coli E* and restimulated with the indicated *E. coli* strains. Student’s t-test. * p <0.05. (F) IFNγ as measured by ELISA from T cells primed with *R*.*gnavus A* and restimulated with bacterial strains from the indicated taxa. (G) IFNγ from T cells primed with *R. gnavus A* and restimulated with two strains of *Ruminococcus*. Student’s t-test. ** p <0.01. (H) IFNγ as measured by ELISA from T cells primed with *B. ovatus C* and restimulated with bacterial strains from the indicated taxa. Student’s t-test. **** p <0.0001. (I) IFNγ from T cells primed with *B. ovatus C* and restimulated with either *B. ovatus A* or *B. ovatus B*. Student’s t-test. * p <0.05. (J) Relationship between the pairwise IFNγ distance relative to the priming strain and the percent whole-genome similarity relative to the priming strain.

We next examined the specificity of T cell responses towards our antigen library using these three *in vitro* primed T cell cultures (Supplementary Table 4). We found that *E. coli-*primed T cells produced more IFNγ in response to *E. coli* strains than to strains belonging to Bacteroides, Firmicutes and Actinobacteria respectively (p < 0.0001, Student’s t-test, Figure 1D). Interestingly, IFNγ secretion was greatest following restimulation with the *E. coli* strain used to prime the T cells (*E. coli E* red dots, Figures 1D and 1E), suggesting that T cell responses towards the priming strain are stronger than towards other strains of *E. coli*.

We also examined the specificity of T cells primed with antigens derived from *R. gnavus* strain *A* and *B. ovatus strain C* (Supplementary Table 4). Restimulation of the *R. gnavus* primed T cells with *R. gnavus* elicited the greatest overall response (red dot, Figure 1F). We found that IFNγ responses to the priming *R. gnavus* strain were stronger than responses towards the other *Ruminococcus* strains in our library (p <0.01, Student’s t-test, Figure 1G), confirming that T cells primed *in vitro* by a single strain have specificity towards the priming strain. We also found that *B. ovatus*-specific T cells produced more IFNγ in response to *B. ovatus* antigens compared to antigens from strains of Firmicutes or Actinobacteria (p < 0.0001, Student’s t-test, Figure 1H). In addition, the *B. ovatus* strain used for priming elicited the greatest overall response amongst the other *B. ovatus* strains in our library (*B. ovatus strain C*, Figure 1I). Of note, we found the response of *B. ovatus* primed T cells towards Proteobacteria to be as strong as responses towards the priming antigen (Figure 1F), which indicates that non-specific, perhaps LPS driven, responses are also captured in this model system.

We also considered whether the magnitude of T cell responses correlated with the genomic similarity relative to the priming strain. We compared the whole-genome similarity of *E. coli* strains in our set relative to the priming strain (*E. coli strain E*) and found that T cell responses correlated with total genomic similarity across strains of *E. coli* (R^2^ = 0.91, p = 0.007, Figure 1J). IFNγ T cell responses also correlated with total genomic similarity across strains of *Ruminococcus* (R^2^ = 1.0, p = 0.02, Figure 1J) and across strains of *B. ovatus* (R^2^ = 0.86, p = 0.17, Figure 1J), suggesting the genomic similarity across related bacterial strains associates with the degree to which CD4^+^ T cells can respond to antigens from these strains.

These data collectively demonstrate that T cells primed *in vitro* with single bacterial strains are useful for interrogating T cell specificities at scale. T cells primed *in vitr*o with antigens from microbiota-derived strains accurately distinguish between microbiota-derived antigens derived from different phylum and can distinguish between strains of the same species, consistent with what is described for gut microbiota-specific antibodies (Yang et al., 2022). Lastly, relative genome similarity between related strains explains the magnitude of T cell responses across related strains.

### Lamina propria primed Th17 cells distinguish between gut-derived microbial strains

To extend our *in vitro* observations to an *in vivo* model, we considered whether T cells in the lamina propria respond specifically to microbiota-derived antigens. Several studies have characterized individual gut-microbial strains that induce mucosal Th17 cells (Atarashi et al., 2015; Britton et al., 2020; Ivanov et al., 2009; Tan et al., 2016; Viladomiu et al., 2017). Furthermore, antigen-specific Th17-skewed responses to the model organism segmented filamentous bacteria (SFB) are well-described (Yang et al., 2014). We applied the strategies used to delineate cognate responses towards organisms such as SFB to define CD4^+^ T cell specificities towards human microbiota-derived commensal strains. To establish these methods, we colonized germ free C57Bl/6 mice with microbiota containing SFB and two weeks later purified T cells from the lamina propria. We restimulated these lamina propria T cells *ex vivo* with APC loaded with antigens from a stool lysate containing SFB and measured T cell responses using an IL-17A ELISpot. As measured by the number of spot forming cells (SFC) by ELISpot, we found that IL-17A-secreting cells isolated from animals colonized with SFB responded more strongly to lysates containing SFB compared to SPF fecal lysates not containing SFB (p = 0.03, Student’s t-test), consistent with what has been previously described (Yang et al., 2014). To expand the scale of our experiments, we confirmed that responses from T cells isolated from mesenteric lymph nodes (mLN) correlated with responses from T cells isolated from the lamina propria of the same animal (R^2^ = 0.63, p = 6.4×10^−6^, Pearson correlation, Figure S2A).

Having established a methodology for measuring Th17 cell specificities *ex vivo*, we examined specificities towards human microbiota-derived bacterial strains. To explore *in vivo* educated T cell specificities, we monocolonized germ-free animals with one of six bacterial strains isolated from a human donor (Supplementary Tables 1 and 5) and measured IL-17A secretion by single T cells following restimulation using an ELISpot assay (Figure 2A). *Ex vivo* IL-17A responses from T cells isolated from germ-free animals were lower in response to all six strains compared to colonized animals (Figure 2B). For most strains, T cell responses towards the colonizing strain were higher than responses to the other strains in the set (Figures 2B and S2B). These results demonstrate the Th17 cells which develop following colonization with a single bacterial stain have a degree of specificity towards that strain, but some cross-reactivity between species also occurs.

**Figure 2.**
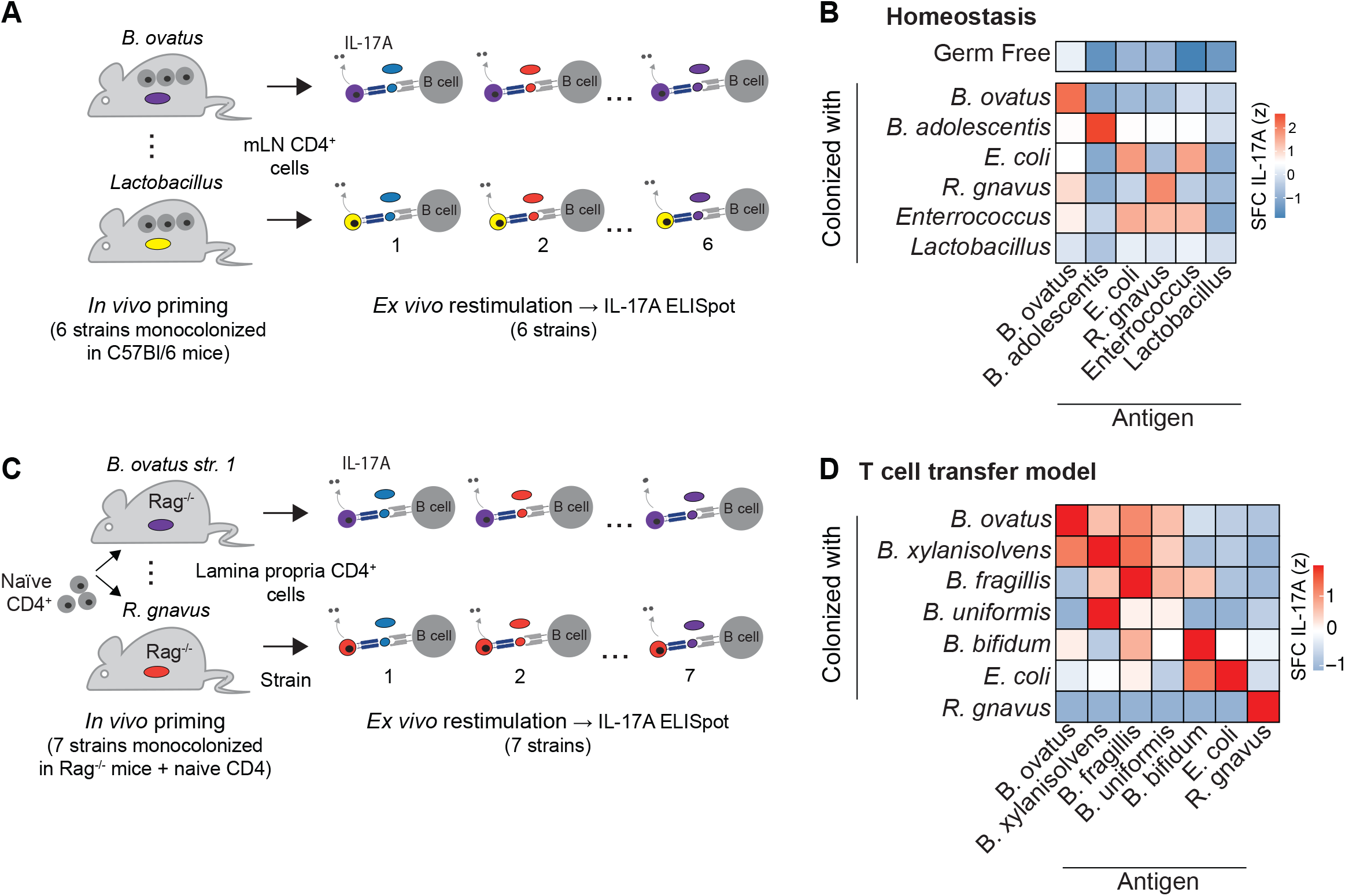
Lamina propria Th17 cells distinguish between gut-derived microbial strains. (A) Schematic of experimental design. 50,000 CD4^+^ T cells from the mLN of ex-germ free mice colonized with a single bacterial strain were restimulated *ex vivo* with the indicated lysates using CD11c^+^ dendritic cells (DC) as antigen presenting cells. T cell reactivity was measured using an IL-17A EliSPOT. (B) Germ-free were colonized with the indicated bacterial strains for two weeks and CD4^+^ T cells from the mLN were restimualted *ex. vivo* with the indicated strains. IL-17A was measured by EliSPOT. SFC, spot forming cell. (C) Schematic of experimental design. 1×10^6^ naïve CD4^+^ CD45RB^hi^ T cells were transferred into Rag^-/-^ mice colonized with a single bacterial strain. Seven weeks later, T cells from the lamina propria were restimulated *ex vivo* with the indicated lysates using CD11c+ DCs as antigen presenting cells. T cell reactivity was measured using an IL-17A EliSPOT. (D) Germ-free Rag^-/-^ mice were colonized with the indicated bacterial strains for two weeks and naïve T cells were adoptively transferred. Lamina propria T cells were isolated seven weeks later and restimulated with the indicated strains. SFC, spot forming cell.

We also assessed the specificity of *in vivo*-primed microbiota-specific T cells in the context of an adoptive transfer model of colitis. We hypothesized the resolution of T cell specificities towards gut microbes would be enhanced in an adoptive transfer model since: 1) the size of the T cell repertoire would be constricted compared to the repertoire of a wildtype mouse and 2) microbiota-specific T cells would be enriched in the absence of endogenous host T cells. We included an additional four strains of *Bacteroides* and two unique strains of *B. ovatus* to better examine how lamina propria T cells distinguish between more-closely related strains (Supplementary Tables 1 and 6). Using this approach, we monocolonized Rag1^-/-^ mice with individual bacterial strains, transferred 1×10^6^ naïve T cells and seven weeks later interrogated the specificity of lamina propria Th17 cells *ex vivo* as before (Figure 2C). Mice colonized with individual bacterial strains did not develop overt colitis, in contrast to mice colonized with all seven strains as a community (Figure S2C). As observed in the wildtype setting, lamina propria T cell responses from monocolonized mice were greatest towards the colonizing strain (Figures 2D and S2D). Of note, we found mucosal Th17 responses towards *R. gnavus* to be highly specific (Figures 2D and S2D). This stands in contrast to the mouse Ig receptor which binds certain *R. gnavus* strains non-specifically due to the expression of a B-cell superantigen (Bunker et al., 2019; Yang et al., 2022).

We examined the relationship between the magnitude of T cell responses and the pairwise distance in the 16S gene sequence between strains. Across comparisons, the pairwise 16S similarity correlated with IL-17A T cell responses across the strains in our set (R^2^ = 0.38, p = 5.4×10^−7^, Pearson correlation, Figure S2E). The largest degree of T cell cross-reactivity occurred between more closely related strains from genus *Bacteroides* (Figures 2D and S2E), suggesting the magnitude of T cell responses broadly associates with the relative taxonomic distance to the educating strain.

Together, these data demonstrate that T cells primed with antigen from one strain respond more strongly towards closely related strains following restimulation, which suggests that dominant epitopes in the strains studied are unlikely shared between distantly related species. On the other hand, we also find some degree of species-level specificity that develops in the pool of Th17 cells following colonization with strains of *Bacteroides*, similar to that of the specificity of IgA towards microbiota-derived strains (Yang et al., 2022).

### *Ex-vivo* T cell restimulations are predictive of Th17 responses *in vivo*

Our *ex vivo* T cell restimulation approach using individual strains was specific enough to detect differences at the species or strain level (Figure 2D), therefore we considered whether this approach might also distinguish the relative contribution of individual strains to the total microbiota-specific T cell pool that develops in mice colonized with more complex communities. To explore these questions, we colonized germ free mice with defined communities representing the culturable fraction of an individual’s stool sample (Faith et al., 2014, 2010; Goodman et al., 2011). We found that these communities elicit varying proportions of RORγt^+^ FoxP3^-^ CD4^+^ T cells (Th17 cells) within the colonic lamina propria (p = 0.0005, ANOVA, Figure 3A), consistent with our previous findings (Britton et al., 2019).

**Figure 3.**
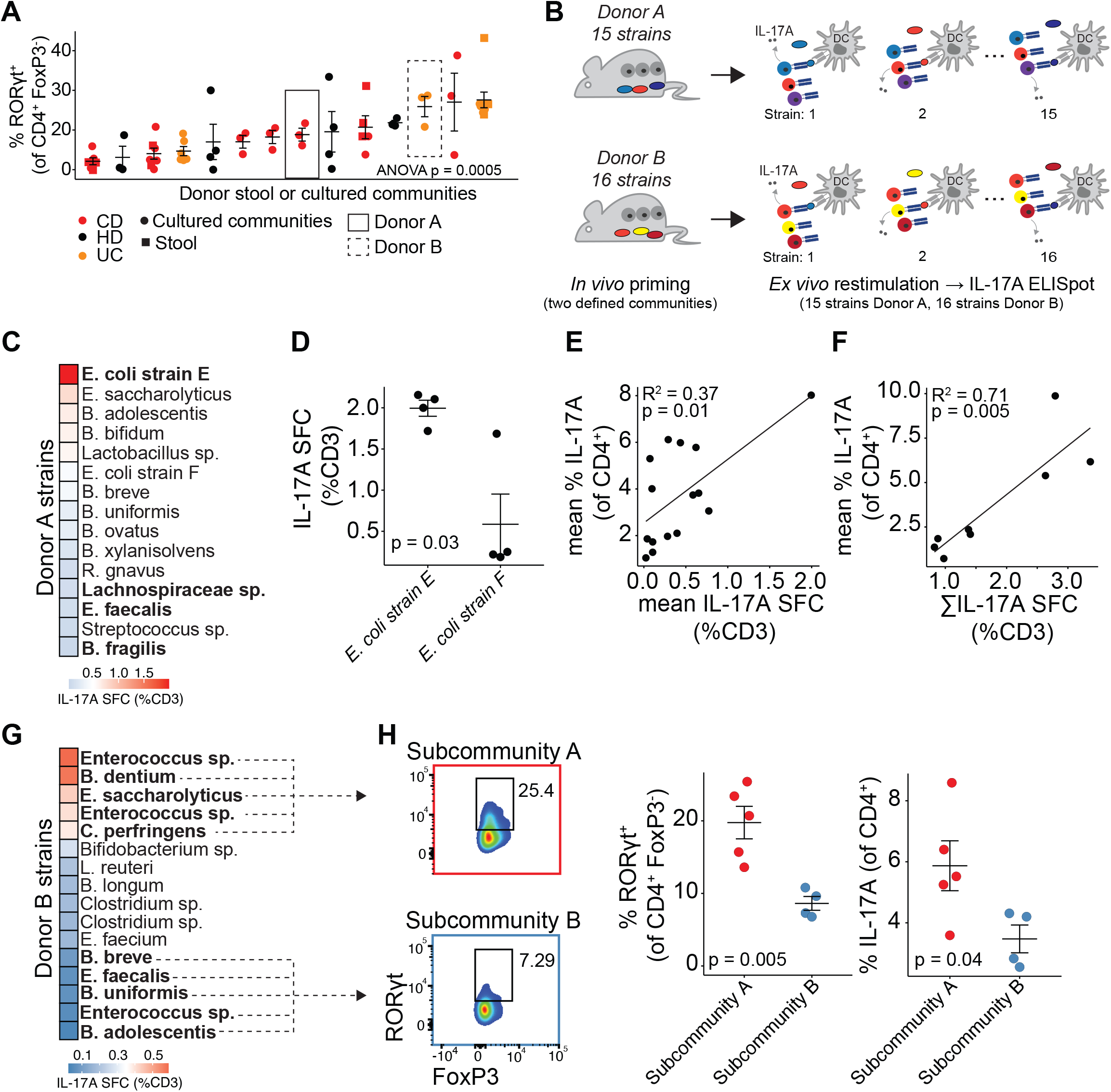
*Ex-vivo* T cell restimulations are predictive of Th17 responses *in vivo*. (A) Germ-free mice were colonized with either whole stool or defined culture communities derived from different donors and colonic T cells were analyzed by flow cytometry. Donor A and Donor B are highlighted. (B) Schema of experimental design. Germ free mice were colonized with defined culture communities. Two weeks later CD4^+^ T cells were isolated from the mLN and co-cultured with CD11c^+^ DC’s loaded with antigen from the individual bacterial strains contained within the defined culture community. T cells were assayed using an IL-17A ELISPOT. (C) CD4^+^ T cells from mice colonized with Donor A’s culture community were restimulated with individual bacterial strains. T cell responses were measured via IL-17A ELISPOT 48 hours after re-stimulation. SFC, spot forming cell. (D) Th17 cells from mice colonized with Donor A’s culture community preferentially respond to *E. coli strain E*. (E-F) Germ-free mice were colonized with orthogonally fractionated subcommunities from Donor A’s culture community and colonic CD4^+^ T cells were analyzed by flow cytometry. (E) Relationship between the average IL-17A response induced by each strain *ex vivo* and the percent colonic CD4^+^ IL-17A^+^ in subcommunities where that strain was present. (F) The average percent colonic CD4^+^ IL-17A^+^ induced by a given subcommunity correlates with the sum of the *ex vivo* IL-17A T cell responses for strains within that subcommunity. (G) CD4^+^ T cells from mice colonized with Donor B’s culture community were restimulated with individual bacterial strains. T cell responses were measured via IL-17A ELISPOT 48 hours after re-stimulation. SFC, spot forming cell. (H) Germ-free mice were colonized with Subcommunity A and Subcommunity B based off the results of Figure 3G and colonic CD4^+^ T cells were analyzed by flow cytometry.

To determine if combining natural *in vivo* priming with *ex vivo* restimulation could identify the strains responsible for modulating the Th17 phenotype induced by a given microbiota, we first colonized mice with a defined community from a donor with Crohn’s disease (Donor A, Figure 3A). We chose Donor A since it includes a Th17-inducing *E. coli* we previously identified using a brute-force combinatorial gnotobiotic screen. The screen was designed to identify dominant effector strains that drive a specific phenotype (Ahern et al., 2014; Britton et al., 2020; Faith et al., 2014). To determine if we could identify cognate Th17 responses towards this strain of *E. coli* using our more efficient *ex vivo* approach, we profiled T cell responses one-at-a-time towards each of the strains contained within the community of Donor A using an IL-17A ELISpot as before (Figure 3B). We found the majority of Th17 cells from the mLN of these mice recognized antigens from a specific *E. coli* strain (*E. coli strain E*, Figures 3C and S3A). *E. coli strain E* is the same strain we previously identified as a potent Th17-inducing effector strain (Britton et al., 2020). Interestingly, T cell responses following restimulation with *E. coli strain E* were greater than responses with a second *E. coli* strain (*E. coli strain F*) also isolated from Donor A (p = 0.03, Student’s t-test, Figure 3D). Therefore, our *ex vivo* restimulation approach correctly identified a dominant effector strain from a complex community using a single batch of 6 germ-free mice compared to the 50 mice used in our original combinatorial strategy (Britton et al., 2020) and differentiated between two strains of *E. coli*.

Previous approaches attempting to identify strains that influence host immune phenotypes have focused on dominant effector strains that have large influences on the *in vivo* immune signature (Britton et al., 2020; Ivanov et al., 2009; Mazmanian et al., 2008; Yang et al., 2022). However, the immune phenotype established by a gut microbiota likely results from a combination of contributions from multiple organisms. We hypothesized the *ex vivo* approach that identified the dominant Th17 *E. coli strain E* might also quantify the smaller contributions from the remaining strains in the community. If these smaller contributions are at least partially additive, we can predict the baseline immune tone from community composition using subsets of the original community. We compared the proportion of antigen-specific IL-17A^+^ cells measured *ex vivo* versus the actual Th17 levels from gnotobiotic mice colonized with orthogonally fractionated subcommunities from Donor A. Fractions containing *E. coli strain E* elicited higher Th17 cells in the colonic lamina propria compared to fractions containing strains eliciting weak T cell responses as measured by ELISpot *ex vivo* (for example, *Lachnospiraceae* p < 0.01, *E. faecalis* p < 0.01 and *B. fragilis* p < 0.03, Student’s t-test, Figure S3B). Our approach also predicted how each strain impacted the average Th17 induction across fractions of Donor A’s culture community (CD4^+^IL-17A^+^ R^2^ = 0.37 p = 0.01 and RORγt^+^ FoxP3^-^ CD4^+^ R^2^ = 0.26 p = 0.03, Pearson correlation, Figures 3E and S3C). Remarkably, by summing the individual *ex vivo* T cell response to the strains in a subcommunity, our approach predicted the Th17 phenotype elicited by each community *in vivo* (CD4^+^IL-17A^+^ R^2^ = 0.71, p = 0.005 and RORγt^+^ FoxP3^-^ CD4^+^ R^2^ = 0.55 p = 0.02, Pearson correlation, Figures 3F and S3D). These data show that the ability of a single strain to induce Th17 cells in the gut under homeostatic conditions is directly linked to the antigenicity of that strain. Furthermore, in the context of small communities, we show an additive relationship between the capacity for strains to restimulate T cells and the proportion of Th17 cells induced within the gut lymphoid tissue.

To determine if this approach generalizes to other human gut microbiota communities, we examined a second microbiota from a donor with ulcerative colitis (Donor B, Figure 3A). As before, we colonized mice with a defined culture community and profiled specific T cell responses towards strains isolated from Donor B one-at-a-time (Figure 3B). We found the pattern of T cell-specificities towards Donor B strains differed from the pattern observed in response to strains from Donor A. Rather than a single strain eliciting a dominant response (Donor A, Figure 3C), we found strong Th17 recall responses following restimulation with five strains from the community (Figures 3G and S3E). This included three isolates of genus *Enterococcus*, and two other more distantly related species. Interestingly, three additional distinct *Enterococcus* isolates did not generate an IL-17A response in the restimulation assay, again demonstrating the specificity by which the Th17 repertoire is shaped by individual strains. Notably, the culture community from Donor B did not contain an *E. coli* strain.

We next colonized gnotobiotic mice with subcommunities comprising either the five most antigenic strains or five non-antigenic strains from the Donor B community (Subcommunity A and Subcommunity B respectively, Figure 3H). These communities shared similar phylogenetic composition. As with Donor A, the measured antigenicity of the strains was an excellent predictor of the Th17-inducing capacity of those strains. Mice colonized with Subcommunity A (containing strains eliciting high responses) induced more Th17 cells in the colonic lamina propria compared to mice colonized with Subcommunity B (p = 0.005 and p = 0.04 respectively, Student’s t-test, Figure 3H). Of note, we did not observe a difference in the fraction of RORγt^+^ FoxP3^+^ CD4^+^ (RORγt^+^ Treg) cells induced in response to colonization between Subcommunity A and Subcommunity B (Figure S3F). Collectively, these results demonstrate that *ex vivo* T cell restimulations quantified by ELISpot efficiently identify the *in vivo* contributions of effector strains within complex communities.

The Th17 TCR repertoire specific for SFB is known to bias for Vβ14 usage (Yang et al., 2014). We tested whether the commensal strains contained within our culture libraries also elicit biases in Vβ usage. We colonized mice with either Donor A or Donor B and compared Vβ usage in different tissues within various CD4^+^ T cell subsets by spectral flow cytometry. Mice colonized with SFB and germ-free mice were included as controls (Supplementary Table 7). We found that Vβ usage broadly segregated the dataset into lymphoid-derived and non-lymphoid derived tissues (Figure S3G). We did not find enrichments or consistent Vβ usage within any tissue following colonization with either Donor A or Donor B. We did confirm, however, that Th17 cells within the small intestine of mice colonized with SFB or sourced from facilities containing SFB (Taconic Farms) contain a bias in Vβ14 usage (Figure S3H).

### T cell specificities for microbiota-derived strains differ during homeostasis versus inflammatory settings

The success predicting Th17 phenotypes during homeostasis using *ex vivo* restimulations led us to consider whether these methods might predict T cell responses during colitis. To test this idea, we colonized Rag1^-/-^ mice with culture communities from Donor A and Donor B and adoptively transferred naïve T cells as previously described (Figure 4A) (Britton et al., 2019). Naïve T cells transferred into mice colonized with communities from Donor A or Donor B resulted in colitis as measured by weight loss (Figure S4A). Six weeks after transfer, we isolated T cells from the lamina propria and profiled T cell responses using an IL-17A ELISpot as before (Figure 4A). Surprisingly, we found that Th17 responses towards strains within Donor A’s culture community were biased towards species of *Bacteroides* six weeks after adoptive transfer (p <0.01, Student’s t-test, Figure 4B). Using a similar experimental design, we also examined the antigen-specific response in reconstituted Rag1^-/-^ mice colonized with the defined community from Donor B. As with Donor A, we found that a broad range of strains stimulated CD4^+^ T cells from these colitic mice and that the greatest IL-17A recall response was to a strain of *Bacteroides* (Figures 4B and 4C). Importantly, Th17 cell responses to strains within Donor A’s culture community were similar across time and across separate batches of mice demonstrating that the specificity to *Bacteroides* during colitis is not because of a stochastic skewing of the naïve CD4 T cell repertoire in one experiment (R^2^ = 0.74, p = 5.6×10^−6^, Pearson correlation, Figure S4B). Th17 cells that co-secrete IL-17A and IFNγ are particularly implicated in inflammatory intestinal responses (Ahern et al., 2010). We found using a FluoroSPOT assay that strains eliciting IL-17A SFC correlated with strains eliciting IFNγ SFC at six-weeks post-transfer (R^2^ = 0.65, p = 7.5×10^−9^, Pearson correlation, Figure S4C) in mice colonized with Donor A.

**Figure 4.**
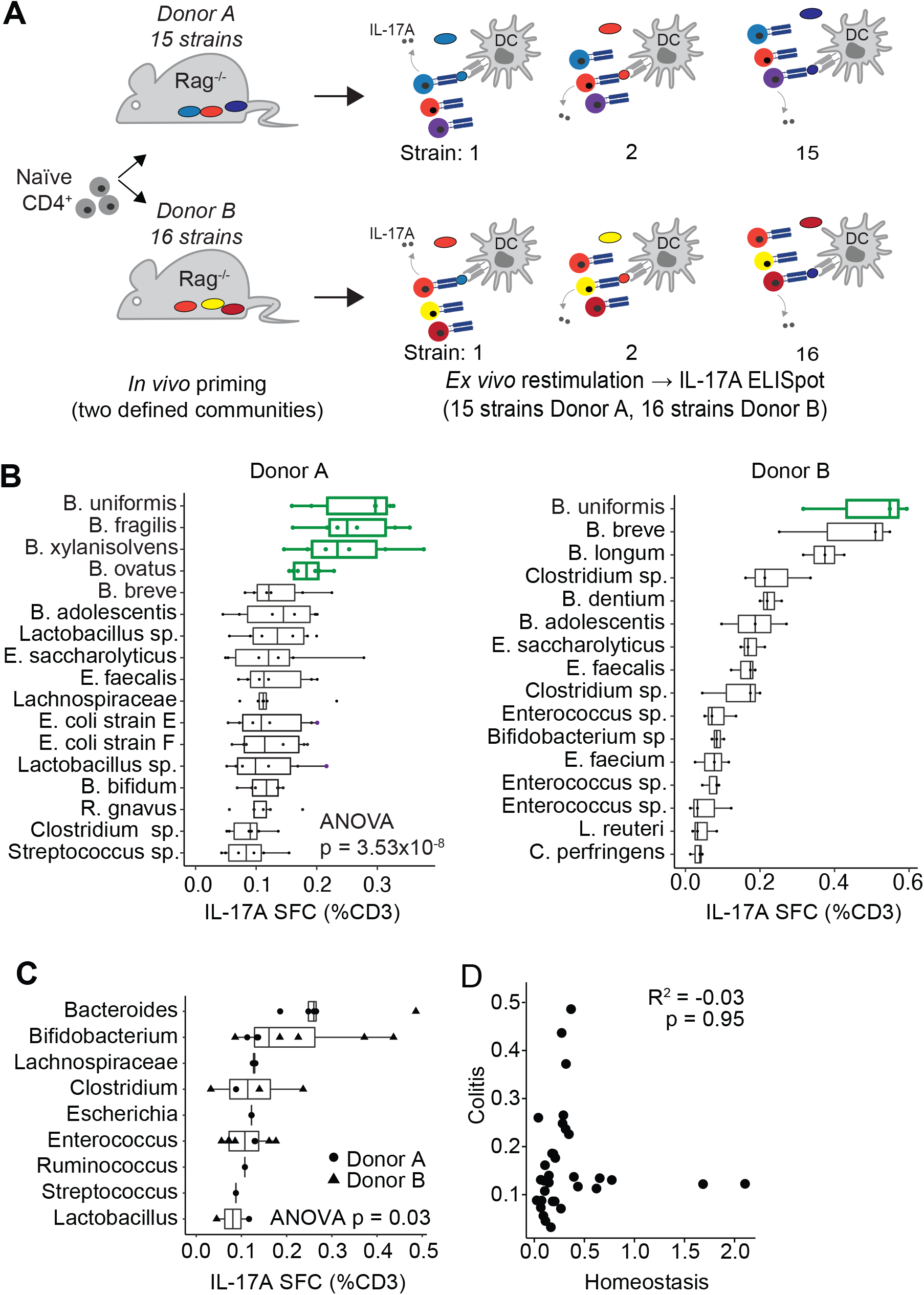
T cell specificities for microbiota-derived strains differ during homeostasis versus inflammatory settings. (A) Schema of experimental design. Rag^-/-^ mice were colonized with defined culture communities for two weeks and 1×10^6^ naïve T cells were adoptively transferred. Lamina propria T cells were isolated six weeks later and restimulated with individual bacterial strains using CD11c^+^ DC’s to present antigen. (B) Lamina propria CD4^+^ T cells isolated from mice colonized with cultured communities from either Donor A or Donor B were restimulated with the indicated. T cell responses were measured via IL-17A ELISPOT 48 hours after re-stimulation. SFC, spot forming cell. Members of *Bacteroides* are colored in green. (C) IL-17A SFC in response to restimulations from strains in the indicated taxa across both Donor A and Donor B cultured communities. (D) IL-17A SFC in response to stimulation from strains in both Donor A and Donor B in homeostatic versus colitis settings.

Lastly, we compared IL-17A T cell responses to individual strains across both Donor A and Donor B in conditions of homeostasis (wildtype C57Bl/6J mice) and in Rag^-/-^ mice following adoptive transfer colitis. There was no correlation in the T cell reactivities towards specific strains across homeostatic and colitis settings (R^2^ = -0.03, p =.95, Pearson correlation, Figure 4D), demonstrating that even in the context of identical community composition, T cells specificities are enriched towards different bacterial strains during homeostasis as compared to during inflammation.

## DISCUSSION

There is great interest in determining how specific strains within the gut microbiome modulate the host immune system with multiple potential applications for treating or preventing immune mediated disease. Identifying immunogenic bacterial strains in the setting of a complex microbiota and determining the resolution of immune-microbiota interactions remain key challenges. Here, we use a combination of anaerobic culturing, *in vitro* T cell assays, gnotobiotic mouse models, and *ex vivo* T cell restimulations to explore the influence of species and strain diversity on the specificity of mucosal T cells and the induction of Th17 immunity. Using an *in vitro* system and two different mouse models, we find that mucosal Th17 cells are specific to microbiota-derived strains at the species level and present evidence that microbiota-specific T cells can potentially distinguish between strains of the same species. Our approach correctly identifies the Th17 inducing ability of strains across two complex human gut microbiotas and across both homeostatic and colitis-susceptible models. We demonstrate that this *ex vivo* strain-specific Th17 inducing ability can be used to design novel subsets with predictable influence on the baseline immune tone *in vivo*. To our surprise, we find no correlation between the strains that induce Th17 immunity during homeostasis and the strains that elicit Th17 responses in an adoptive transfer model of colitis. Collectively, our approach represents an efficient discovery tool for parsing immune phenotypes from complex microbiotas and raises the possibility of forward engineering host immune phenotypes using selective microbiota-derived bacterial strains.

Symbiotic microorganisms in the GI tract induce pathology under certain conditions. Microbial species such as *Helicobacter* and SFB are well characterized pathobionts that are known to both potentiate inflammation and also colonize healthy organisms (Chow and Mazmanian, 2010; Stepankova et al., 2007). In addition, stools from human IBD gut microbiotas induce more severe colitis in susceptible mice compared to healthy donor microbiotas despite no discernable differences in terms of composition and diversity (Britton et al., 2019). Determining how microbial organisms exist harmoniously in some contexts but increase the risk for inflammation in others is a fundamental question for the microbiome field. We find that Th17 cells respond to different elements of the microbiota depending on the context, which suggests that 1) changes in the epithelial barrier during inflammation provides bacterial strains differential access to the immune system relative to the conditions during homeostasis or 2) changes in the regulation of the immune system potentiate changes in response to the same collection of bacteria. Towards the later point, studies using TCR transgenic models have shown that commensal-specific T cell responses are restrained during homeostasis but expand in genetically predisposed settings (Xu et al., 2018).

It is unclear if the Th17 microbe-specific responses that we measure are pathogenic. IL-17A is an important cytokine for maintaining epithelial barrier integrity (Lee et al., 2015) and blocking IL-17A can exacerbate IBD (Hueber et al., 2012). Some studies have suggested that IL-17 cellular programs are context dependent and that specific molecules can promote the differentiation of more pathogenic Th17 cells during inflammation (Ahern et al., 2010; Lee et al., 2020). Studies have shown that IFNγ producing T cells are pathogenic in adoptive T cell transfer models (Harbour et al., 2015) and that blocking IFNγ can prevent adoptive transfer colitis (Powrie et al., 1994). Clearly more work is needed to determine which T cell reactivities lead to the induction of colitis and whether the colitogenic potential of the microbiota can be efficiently parsed.

The specificity of mucosal T cells towards microbiota-derived antigens is not well-understood. Although antigen-specific interactions between gut-derived bacterial strains and mucosal Th17 cells have been described (Britton et al., 2020; Goto et al., 2014; Ivanov et al., 2009; Viladomiu et al., 2017; Xu et al., 2018), it is not clear how microbe-specific T cells respond to antigens across the phylogenetic space of the microbiome. A recent study found that multiple TCR clonotypes sampled against a defined microbial community react against an epitope widely conserved among Firmicutes (Nagashima et al., 2022). We observe that T cells primed with some microbial strains can similarly cross-react with related species. Notably, we find the strongest T cell responses upon restimulation are generally towards the priming strain, which suggests that some fraction of microbe-specific T cells distinguish between strains of the same species.

Together, these studies suggest that both poly-specific and species-specific T cells exist towards antigens in the gut microbiota. Future work will need to determine the physiologic relevance of highly specific versus more broadly specific T cell reactivities.

Our work has several important limitations. First, our investigation into the specificity of T cells relies upon monocolonizations, which presents the possibility that strains colonize in unnatural anatomic locations and bias immune responses. We recognize the limitations of using monocolonized animals and utilize this model system only as a source to generate microbial-strain specific T cells. Second, our donor-derived culture communities consist of only 15 unique strains. The utility of *ex vivo* restimulation assays to predict immune phenotypes may be more limited in the setting of larger community sizes where the chance of overlapping functionality is greater. Lastly, we focus primarily on IL-17A, which we find to be strongly correlated with IFNg, but we do not examine other important T cell-derived cytokines such as IL-4, IL-10 and IL-22, nor do we examine the specificity of regulatory T cells.

In summary, we describe a method to quickly identify Th17 inducing effector strains from complex bacterial communities using *ex vivo* restimulations. Using an adoptive transfer model of colitis, we find that lamina propria Th17 cells respond to different bacterial strains in homeostatic versus inflammatory conditions. Lastly, we find that T cell responses to microbiota-derived strains are highly specific at the species level, that T cells can differentiate between strains of the same species, and that strain-specific immune responses *ex vivo* predict the baseline immune tone of bacterial communities *in vivo*, which collectively supports the potential to forward engineer bacterial communities for promoting health or preventing disease.

## METHODS

### Recall assay using *in vitro* primed microbiota-specific T cells

*In vitro* recall experiments were performed as previously described with some minor modifications (Gao et al., 2020). X-VIVO15 serum-free media (Lonza) was used to avoid T cell activity to bovine serum proteins. Spleens were harvested from 8–12 week old SPF C57BL/6J mice. Naive CD4^+^ T cells (CD62L^+^ CD44^-^) were purified (Miltenyi Biotec) using an AutoMACS. Splenic DCs (CD11c^+^) were purified using positive selection microbeads (Miltenyi) according to the manufacturer’s instructions. CD11c^+^ DCs were pulsed overnight with bacterial lysates killed via autoclave (1:100 dilution of the stationary phase culture). DC’s and T cells were cocultured at a 1:5 ratio (100,000 T cells/well) for 10 days to prime T cell cultures. On day 10, wells were pooled, washed twice and rested in a 24-well plate for 48hours with 10U/mL IL-2 (R & D systems) to allow for cytokine production to wane. On day 11, B cells (CD19^+^) were purified (Miltenyi) from SPF C57BL/6J mouse spleens and pulsed overnight with a panel of antigens derived from 35 unique bacterial strains (1:100 dilution). On day 12, loaded B cells were irradiated (12Gy) using an x-ray irradiator, washed twice and cocultured with rested T cells at a 2:1 B to T cell ratio. Coculture supernatants were collected 48 hours later and analyzed by ELISA (R&D Systems).

### Mice and gnotobiotic experiments

All animal studies were carried out in accordance with protocols approved by the Institutional Animal Care and Use Committee in Icahn School of Medicine at Mount Sinai. For adoptive transfer experiments, six to eight-week-old WT C57BL/6J (Jackson Laboratory stock #000664) mice were purchased from Jackson Labs. Mice were kept under specific pathogen-free conditions at the Icahn School of Medicine at Mount Sinai Gnotobiotic Facility. Germ free C57BL/6J and Rag1^−/−^ C57BL/6J were bred in isolators at the Icahn School of Medicine at Mount Sinai Gnotobiotic Facility. Mice were colonized by oral gavage with either pure cultures of isolated strains or a pooled cultured community at 6-8 weeks old. Following colonization, gnotobiotic mice were housed in barrier cages under positive pressure and handled using strict aseptic technique. Mice were colonized for 2 weeks before analysis of lamina propria cell populations. Some gnotobiotic experiments were performed as a part of other published studies to understand relationships with mucosal immune phenotypes (Britton et al., 2020).

### Bacterial culturing and donor community construction

Stool samples from Donor A (Crohn ‘s disease) and Donor B (UC) were collected from de-identified donors in remission and were quickly frozen following donation before processing. Arrayed culture communities from donor stool samples were generated as previously described (Britton et al., 2020; Faith et al., 2010; Goodman et al., 2011). Briefly, a variety of solid medias were inoculated with a fecal slurry and incubated under anaerobic and aerobic conditions.

Colonies were inoculated into liquid media in anaerobic conditions and arrayed into multi-well plates (LYBHIv4 (46); 37 g/L Brain Heart Infusion [Becton Dickinson], 5 g/L yeast extract [Becton Dickinson], 1 g/L each of D-xylose, D-fructose, D-galactose, cellubiose, maltose, sucrose, 0.5 g/L N-acetylglucosamine, 0.5 g/L L-arabinose, 0.5 g/L L-cysteine, 1 g/L malic acid, 2 g/L sodium sulfate, 0.05% Tween 80, 20 μg/mL menadione, 5 mg/L hemin [as histidine-hemitin], 0.1 M Mops, pH 7.2). Strains comprising each donor culture community were characterized by a combination of matrix assisted laser desorption/ionization time-of-flight mass spectrometry (MALDI-TOF), 16S ribosomal DNA amplicon and whole-genome shotgun sequencing. Culture communities were kept arrayed for *ex vivo* recall experiments or pooled as a cocktail for administration to mice.

### Lamina propria lymphocyte isolation

Gut tissues were prepared as previously described (Britton et al., 2019). Briefly, cleaned gut tissues were deepithelialized in 5 mM ethylenediaminetetraacetic acid (EDTA), 15 mM Hepes, and 5% fetal bovine serum (FBS) in Hank’s Balanced Salt Solution (HBSS) before digestion with 0.5 mg/mL Collagenase Type IV (Sigma Aldrich) and 0.25 mg/mL DNase 1 in HBSS with 2% FBS. Lymphocytes enriched by passing the cell suspension sequentially through 100- and 40-μm strainers. For enzyme-linked immune absorbent spot (ELISpot) experiments, samples were passed through a Dead Cell Removal Kit (Miltentyi Biotech) to remove debris.

### Flow cytometry

Flow cytometry of CD4^+^ T cells was performed as previously described (Britton et al., 2019). The following antibodies were used for T cell phenotyping: CD45 APC-Cy7 (Biolegend), CD4 APC (Biolegend), FoxP3 PE (Thermo Fisher/ eBioscience), RORγt PerCp-Cy5.5 (BD Bioscience), and IL-17A-PE (Biolegend), and dead cells were excluded using Zombie NIR (Biolegend). To detect intracellular IL-17A, isolated lymphocytes were first restimulated with 5 ng/mL phorbal 12-myristate 13-acetate and 500 ng/mL ionomycin in the presence of monensin (Biolegend) for 3.5 h. FoxP3 and RORγt expression was analyzed in unstimulated cells. The Vβ chain repertoire analysis used the following antibodies: Vβ2 AF647 (Biolegend), Vβ3 BrilliantViolet 510 (BD), Vβ5.1/5.2 PE-Cy7 (Biolegend), Vβ6 BrilliantViolet 650 (BD), Vβ7 FITC (Biolegend), Vβ8.1/8.2 APC-Vio770 (Miltenyi), Vβ10b BrilliantViolet 711 (BD), Vβ11 BrilliantViolet 421 (BD), Vβ12 BrilliantViolet 480 (BD), Vβ13 PerCP-eFluor710 (ThermoFisher/eBioscience), Vβ14 biotin (ThermoFisher/eBioscience) with streptavidin Qdot800 (ThermoFisher), Vβ17a BrilliantViolet 605 (BD), CD45 BrilliantViolet 750 (BD), FoxP3 PE (ThermoFisher/eBioscience) RORγt APC (BD) and CD4 BrilliantViolet 570 (Biolegend). T cell phenotyping data and Vβ chain phenotyping data were acquired using an Aurora spectral cytometer (Cytek).

### ELISpot Assay

ELISpot experiments for the detection of microbe-specific T cell responses were performed as previously described with some minor modifications (Yang et al., 2014). For the isolation of pure populations of CD4^+^ T cells, samples were separated by positive selection (CD4^+^ microbeads, Miltenyi Biotech) using an AutoMACS instrument (Miltenyi Biotech). Dendritic cells were isolated from the spleens of naive SPF C57B/6J mice (Jackson Laboratories) by magnetic isolation (CD11c microbeads, Miltenyi Biotech). Bacterial strains were grown to stationary phase in LYBHIv4 media and lysates were prepared by autoclaving the washed bacteria in phosphate-buffered saline (PBS). Presence of SFB was confirmed by PCR. Dendritic cells were plated at a density of 25,000 cells/well and pulsed with antigen (1:100 dilution of the stationary phase culture) overnight. Isolated CD4^+^ T cells were added to the culture the next day (2:1 T cell to dendritic cell ratio) and placed at 37° for 48 h. T cells were generally pooled from 6 mice colonized with the same microbiota in order to obtain sufficient cells to perform these assays. After 48h of coculture, the cells were transferred to an IL-17A ELISpot plate (R&D systems) for an additional 24h and developed according to the manufacturer’s instructions. Experiments using Fluorospot were performed identically but developed according to the manufacturer’s instructions (Immunospot).

### T cell transfer colitis model

Colitis experiments were performed as previously described (Britton et al., 2019). Briefly, CD4^+^ T cells were magnetically enriched (Magnisort CD4 Enrichment Kits; Thermo Fisher) from spleens of naive SPF C57BL/6J (Jackson Labs) before fluorescence-activated cell sorting purification of CD4^+^ CD25^−^ CD45RB^HI^ cells. Rag1^−/−^ C57BL/6 mice were administered 1×10^6^ naive CD4^+^ T cells by intraperitoneal injection and monitored for changes in body mass weekly.

### Comparative bacterial genomics

We calculated genomic distances at the resolution of bacterial strains (Fig. 1J) by comparing the kmer-overlap between each pairwise combination of strains using a kmer size of 20 [cite Faith et.al., biorxiv 2020]. For more distantly related taxa (Fig., S2E), bacterial genomes were annotated by prokka [https://pubmed.ncbi.nlm.nih.gov/24642063/] to extract 16S rRNA sequences with RNAmmer [cite https://pubmed.ncbi.nlm.nih.gov/17452365/]. The 16S rRNA sequences were aligned and pairwise distances were calculated with Clustal Omega [cite https://pubmed.ncbi.nlm.nih.gov/21988835/].

### Statistical analysis and plotting

Analyses throughout this study were performed using RStudio v1.1.463. All p values less than 1×10^−10^ were rounded. Heatmaps were constructed using either *ComplexHeatmap* (Gu et al., 2016) or ggplot2.

## ACKNOWLEDGEMENTS

This work was supported in part by the staff and resources of the Mount Sinai Gnotobiotic Facility, the Mount Sinai Flow Cytometry Core, and the Scientific Computing Division at the Icahn School of Medicine at Mount Sinai. We thank C. Fermin, E. Vazquez, and G.N. Escano for gnotobiotic husbandry and for technical support. We thank Zhihua Li for assistance with bacterial culturing. We thank Dr. Peter Heeger and the members of his laboratory for their help with Fluorospot. This work was supported by National Institutes of Health grants (nos. NIDDK DK112978, NIDDK DK124133, NIDDK DK123749), an NIH F30 to M.P.S. (DK124978), and a SUCCESS philanthropic award and a Crohn’s and Colitis Foundation RFA award to G.J.B. (no. 580924) and J.J.F. (nos. 632758, 651867).

## DATA-SHARING

Whole-genome assembled sequences (FASTA) of strains have been deposited under project number PRJNA637878. All bacterial strains are available upon request.

## AUTHOR CONTRIBUTIONS

M.P.S., G.J.B and J.J.F. wrote the manuscript. M.P.S., P.S., I.M. collected the samples and performed the experiments. M.P.S., G.J.B., J.J.F. analyzed and interpreted the data. All authors read, provided critical feedback and approved the final manuscript.

## FIGURE LEGENDS

**Figure S1.**
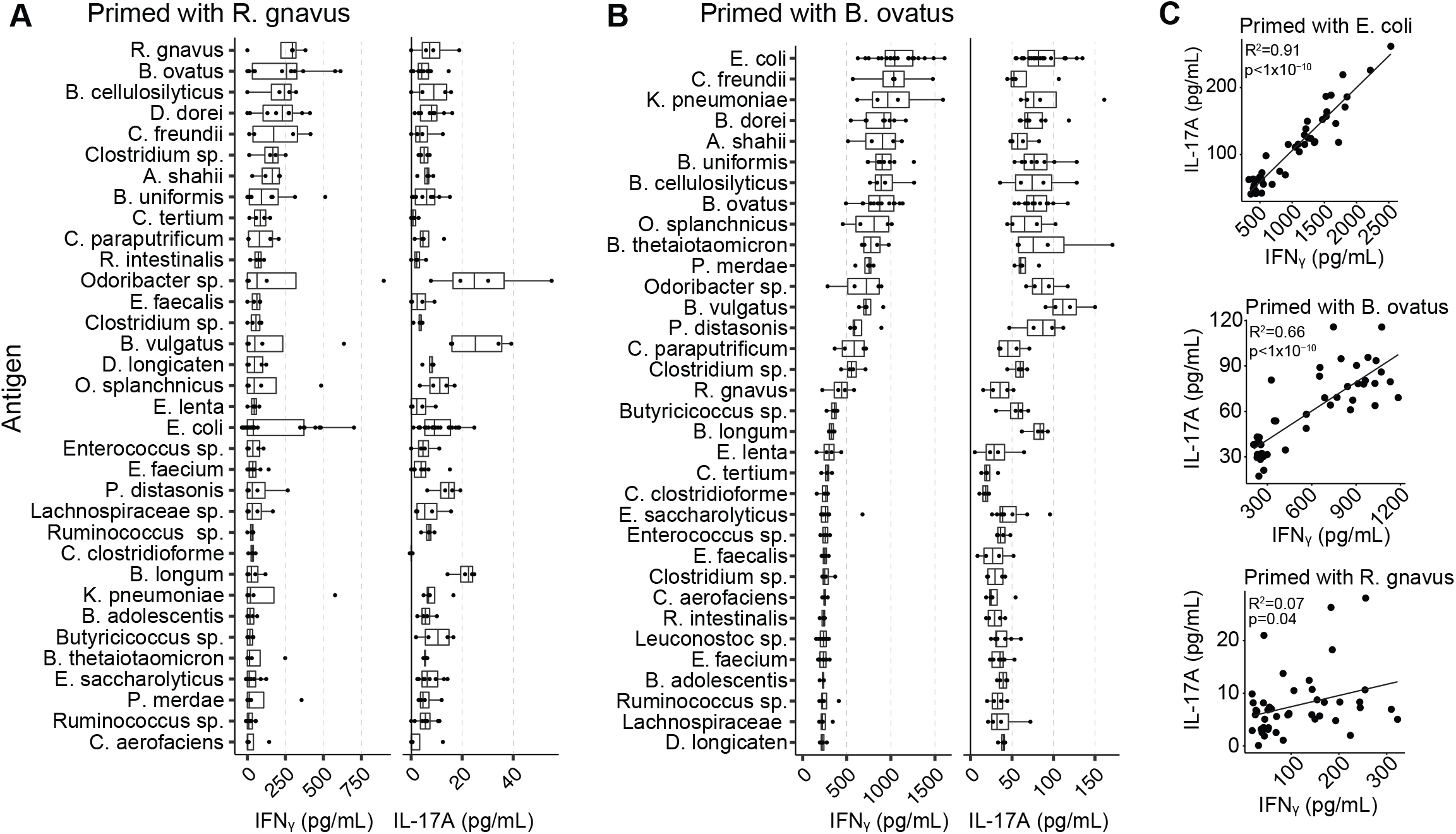
(A and B) IFNγ and IL-17A as measured by ELISA from T cells following restimulation with the indicated bacterial strains. (C) Relationship between IFNγ and IL-17A following restimulation with the 45-member bacterial antigen library.

**Figure S2.**
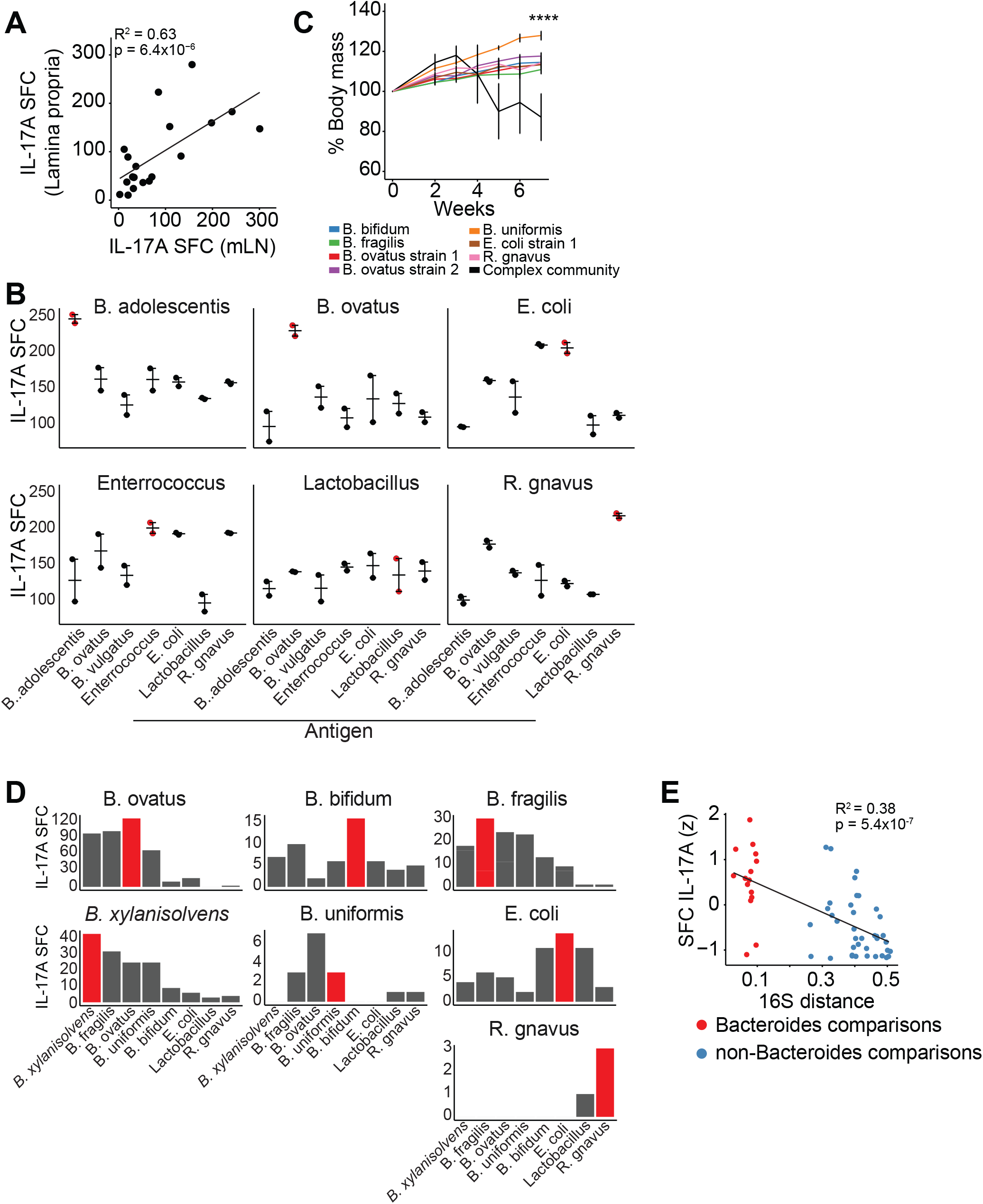
(A) Relationship between SFC of CD4^+^ T cells isolated from the mLN versus the lamina propria. (B) Germ-free were colonized with the indicated bacterial strains for two weeks and CD4^+^ T cells from the mLN were restimualted *ex. vivo* with the indicated bacterial strains. IL-17A was measured by ELISPOT. SFC, spot forming cell. (C) Percent body mass over time following adoptive transfer of naïve CD4^+^ T cells into Rag^-/-^ mice colonized with the indicated bacterial strains. (D) Germ-free Rag^-/-^ mice were colonized with the indicated bacterial strains for two weeks and 1×10^6^ CD4^+^ CD45RB^hi^ naïve T cells were adoptively transferred. Lamina propria T cells were isolated seven weeks later and restimuilated with the indicated strains. SFC, spot forming cell. (E) T cell responses as measured by IL-17A ELISPOT and the 16S distance between the colonizing strain and the restimulation strain. Conditions where the colonizing strain and the restimulation strain are identical were omitted. Strains that elicit a stronger response tend to be more closely related.

**Figure S3.**
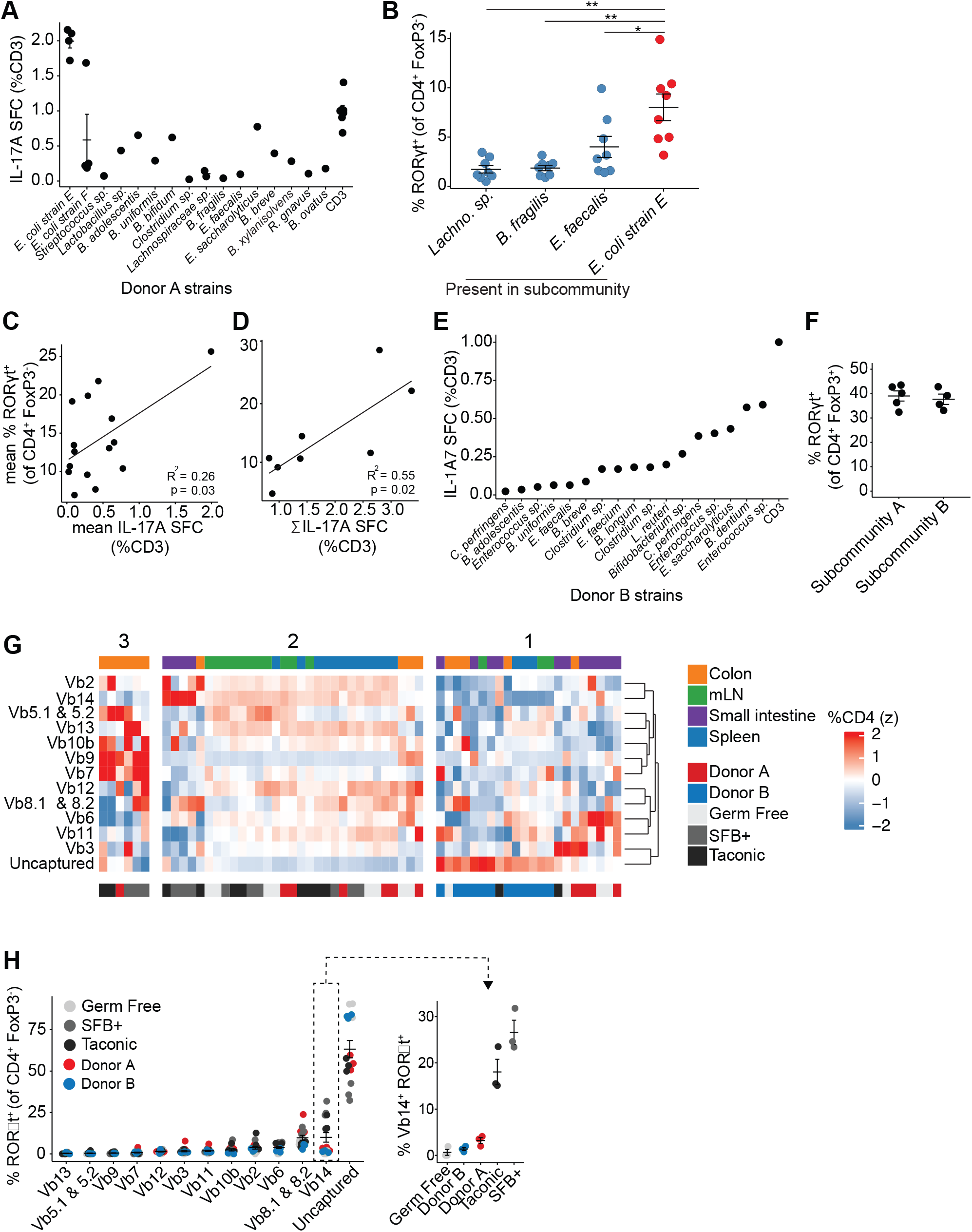
(A) CD4^+^ T cells from mice colonized with Donor A’s culture community were restimulated with indicated conditions. T cell responses were measured via IL-17A ELISPOT 48 hours after re-stimulation. SFC, spot forming cell. (B-D) Germ-free mice were colonized with orthogonally fractionated subcommunities from Donor A’s culture community and colonic CD4^+^ T cells were analyzed by flow cytometry. (C) Th17 induction within the colonic lamina propria by subcommunities where the indicated strain was present. (D) Relationship between the average IL-17A induced by each strain *ex vivo* and the percent Th17 induced *in vivo* by subcommunities where that strain was present. (E) The lamina propria Th17 phenotype induced by a given subcommunity correlates with the sum of the *ex vivo* IL-17A T cell responses for that subcommunity. (E) CD4^+^ T cells from mice colonized with Donor B’s culture community were restimulated with individual bacterial strains. SFC, spot forming cell. (F) Germ-free mice were colonized with colonized with Subcommunity A and Subcommunity B based off the results of Figure 3G and colonic CD4^+^ T cells were analyzed by flow cytometry. (G-H) Mice were colonized with the indicated communities and CD4^+^ T cells were analyzed by spectral flow cytometry for TCR Vβ usage in the indicated tissues.

**Figure S4.**
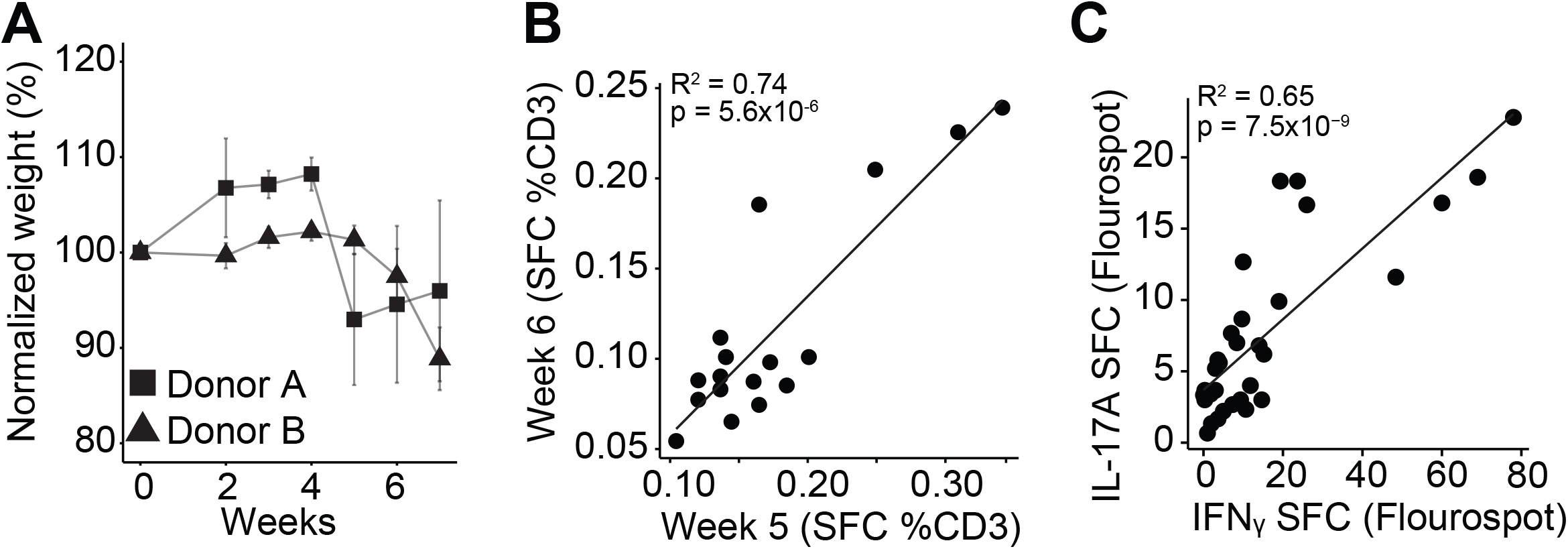
(A) Weight loss over time of Rag^-/-^ mice colonized with Donor A or Donor B following adoptive transfer of 1×10^6^ naïve T cells (B) T cell reactivities towards bacterial strains over time in T cell transfer colitis (C) IL-17A SFC responses correlate with IFNγ responses in T cell transfer colitis

